# When conflict cannot be avoided: executive control dominates early selective sensory modulations during cognitive conflict

**DOI:** 10.1101/177394

**Authors:** Sirawaj Itthipuripat, Sean Deering, John T. Serences

## Abstract

When different sources of sensory information suggest competing behavioral responses, the efficiency of decision-making is impaired. Prior work suggests that at least two mechanisms may play a role in mitigating this interference: using early selective attention to extract the most relevant sensory inputs to avoid conflict or increasing the efficiency of the executive control network to resolve conflict during post-perceptual processing. To test these alternatives, we combined a stimulus-frequency tagging technique with a classic color-word Stroop paradigm, where color-bar targets and letter-string distractors were simultaneously flickered at different frequencies. Using electroencephalography (EEG), we measured the quality of early sensory processing by assessing the amplitude of steady-state visually evoked potentials (SSVEPs) elicited by the targets and distractors. We also measured the engagement of the executive control network by assessing changes in frontal theta (4-7Hz) and posterior alpha oscillations (8-14Hz). Counter to the ‘early selective sensory modulation’ account, the amplitude of the SSVEP response was not modulated by manipulations of color/word congruency, while the frontal theta activity increased and the posterior alpha activity decreased in response to conflict. Moreover, target-related SSVEP amplitude was not correlated with response times (RTs) and a higher (not lower) distractor-related SSVEP amplitude predicted faster RTs. On the other hand, the amplitude of the frontal theta and alpha activity was highly correlated with RTs, irrespective of conflict levels. Over all, these results highlight the dominant role of the executive control network in conflict resolution during post-perceptual processing.

**Significance Statement:** Conflicting information interferes with decision-making. However, this interference can be mitigated either by extracting the most relevant inputs during early sensory processing or by increasing the efficiency of the executive control processes to resolve conflict. By measuring electroencephalography (EEG) in humans performing a modified color-word Stroop task, we examined early sensory responses evoked by targets and distractors while simultaneously monitoring frontal theta and posterior alpha oscillations to index the activation of the executive control network. We found evidence that the executive control network played a more prominent role in resolving conflict.

## Introduction

Distraction caused by the inadvertent processing of task-irrelevant information interferes with the speed and efficiency of decision-making (Jensen and Rohwer JR., 1966; Pashler, 1984; Stroop, 1992; Lavie and Cox, 1997; Wolfe, 1998; Hickey et al., 2010; Anderson et al., 2011; Eckstein, 2011; Awh et al., 2012; Itthipuripat et al., 2015). When task-relevant and task-irrelevant information can be differentiated based on spatial position or low-level features such as orientation or color, early selective attention can facilitate decision-making by modulating gain of sensory responses in visual cortex to bias processing in favor of the relavant stimulus (Moran and Desimone, 1985; Hillyard and Anllo-Vento, 1998; Treue and Martinez-Trujillo, 1999; McAdams and Maunsell, 1999; Reynolds et al., 2000; Martínez-Trujillo and Treue, 2002; Störmer et al., 2009; Scolari et al., 2012; Itthipuripat et al., 2014a, 2014b; Störmer and Alvarez, 2014; Mayo and Maunsell, 2016). Importantly, the magnitude of these early sensory modulations is closely related to behavioral performance (Mangun and Hillyard, 1988; Störmer et al., 2009, 2013; Andersen et al., 2012; Itthipuripat et al., 2013a, 2014a, 2017; Itthipuripat and Serences, 2015; Luo and Maunsell, 2015).

While this ‘early selective sensory processing’ account is supported by data from behavioral tasks that require the processing of low-level sensory features, the extent to which selective sensory processing can support the resolution of cognitive conflict is still in question. For example, in the classic Stroop and Eriksen flanker paradigms, competition arises because different stimuli suggest incompatible semantic interpretations and/or motor plans. In these situations, early selective attention to the task-relevant sensory stimulus might mitigate subsequent post-perceptual conflict by preventing or attenuating any semantic analysis and response planning associated with task-irrelevant stimuli (Appelbaum et al., 2011, 2012; Coste et al., 2011; Zavala et al., 2013). However, contrary to this early selective sensory processing account, other studies have found that conflict interference effects are still observed even when conflicting information is presented at an unattended location or when it is rendered subjectively invisible via visual masking (Nieuwenhuis et al., 2001; Sumner et al., 2007; van Gaal et al., 2008, 2010, 2011; D’Ostilio and Garraux, 2012; Jiang et al., 2015; Padro et al., 2015). This high degree of automaticity suggests that selective sensory processing does not play a substantive role in filtering out conflicting information before it reaches post-perceptual stages. Instead, decision-making efficiency during these tasks may depend primarily upon the activation of the executive control network, including sub-regions of frontal and parietal cortex that support cognitive control functions such as conflict monitoring and task engagement (Carter, 1998; Botvinick et al., 1999, 2001, 2004; Bello et al., 2001; Adleman et al., 2002; Liu et al., 2006; Zimmer et al., 2010; Talsma et al., 2010; Grandjean et al., 2012; Cavanagh and Frank, 2014).

In the present study, we evaluated the relative contributions of the ‘early selective sensory processing’ and the ‘post-perceptual executive control’ mechanisms to decision-making efficiency under high-order cognitive conflict. We combined a stimulus-frequency tagging technique with a classic color-word Stroop paradigm (Jensen and Rohwer, 1966; Stroop, 1992), where task-relevant color-bar targets and task-irrelevant letter-string distractors were flickered at different frequencies (Figure 1a). While human subjects were performing this task, we measured electroencephalography (EEG), which allowed us to examine selective modulations of early visual responses via the simultaneous monitoring of steady-state visually evoked potentials (SSVEPs) elicited by targets and distractors (Norcia et al., 2015). Following many previous studies, we also measured frontal theta activity (4-7Hz) as an index of the activation of the frontal executive control mechanisms (Cavanagh et al., 2011, 2012; Cavanagh and Frank, 2014). Finally, we measured posterior alpha activity (8-14Hz) as an index of the activity of the fronto-parietal network involved in general task engagement (Fries, 2001; Sauseng et al., 2005; Klimesch et al., 2007; Rihs et al., 2007; Fries et al., 2008; Kelly et al., 2009; Zhang et al., 2010; Foxe and Snyder, 2011; Bosman et al., 2012; Sadaghiani et al., 2012; Marshall et al., 2015).

**Figure 1.**
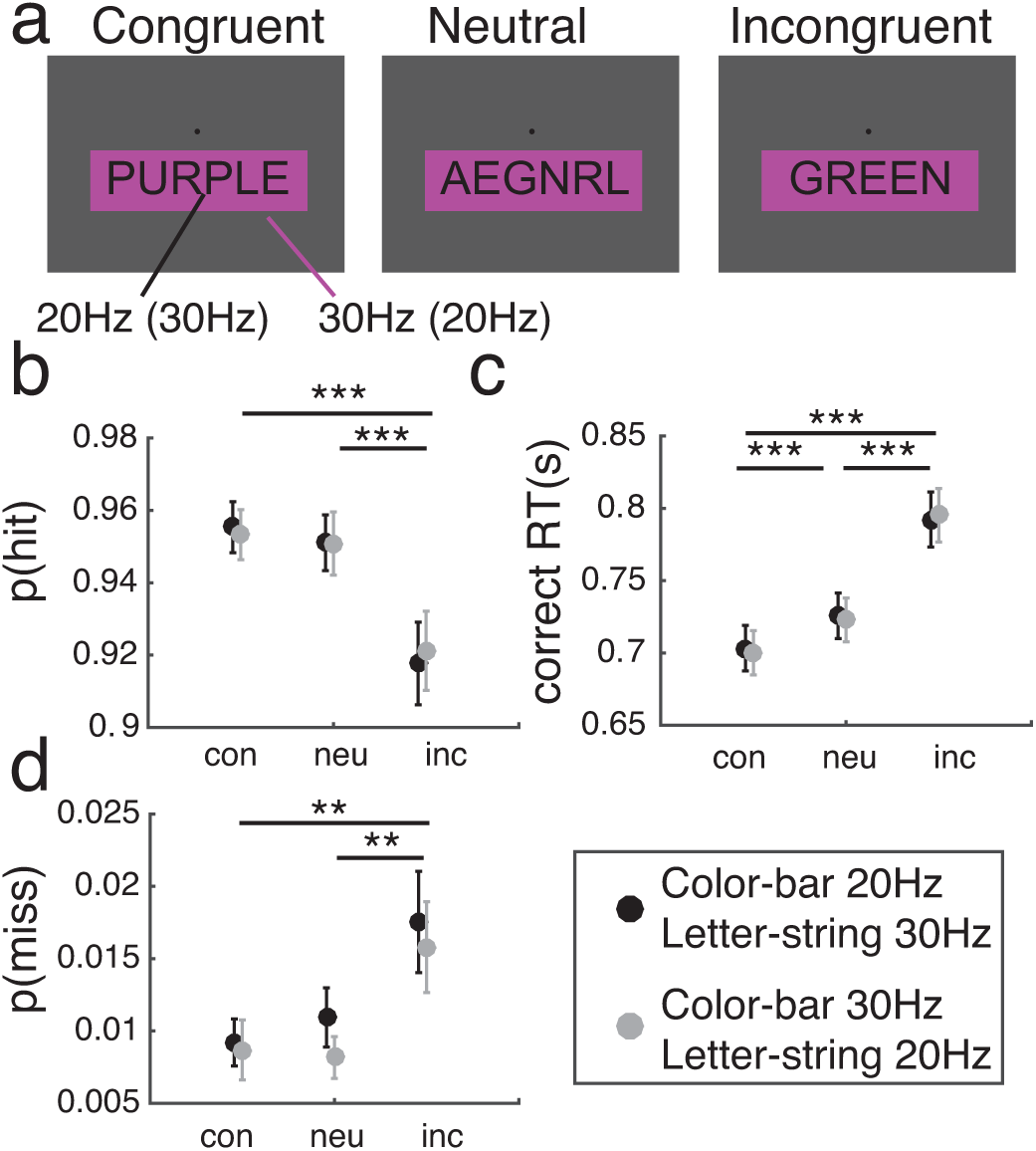
Task design and behavioral results. (a) An adapted version of the classical color-naming Stroop task, where the color-bar (task-relevant target) and the letter-string stimuli (task-irrelevant distractor) were flickered at different frequencies (20Hz and 30Hz, respectively and vice versa). (b) Hit rates, (c) correct RTs, and (d) miss rates differed across congruency conditions but did not differ across blocks where flicker-frequencies assigned to the color-bar and letter-string stimuli were counterbalanced. ** and *** show significant pair-wise differences between congruency conditions with p’s < 0.01 and <0.001. Error bars show ±1 within-subject standard error of mean (S.E.M.).

## Materials and methods

### Subjects

Thirty-one neurologically healthy human volunteers (13 females, 4 left-handed, 18-44 years old) with normal or corrected-to-normal vision were recruited from the University of California, San Diego (UCSD) community. Each volunteer provided written informed consent in accordance with UCSD Institutional Review Board guidelines (IRB#110176), and the experiment was conducted under the protocol that followed the Declaration of Helsinki. Subjects were compensated $15 per hour for participation in the study. Data from one subject were excluded due to excessive EEG blinks, eye and head movement artifacts (>84% of trials), leaving data from 30 subjects in the final behavioral and EEG analyses.

### Stimuli and Experimental Design

Stimuli were presented using MATLAB (Mathworks Inc., Natick, MA) and the Psychophysics Toolbox (version 3.0.8; Brainard, 1997; Pelli, 1997) on a PC running Microsoft Windows XP. Subjects were seated 60 cm from a CRT monitor (with a dark grey background of 4.11 cd/m^2^ ± 0.12 SD, 60Hz refresh rate) in a sound-attenuated and electromagnetically shielded chamber (ETS Lindgren). The entire experiment (EEG preparation, experimental tasks, and breaks) lasted approximately 2-2.5 hours.

Like many previous Stroop studies that required manual responses instead of verbal responses (e.g., Krebs et al., 2010, 2013; Appelbaum et al., 2012; Donohue et al., 2013, 2016; van den Berg et al., 2014), subjects first underwent a stimulus-response mapping task in which they learned to associate the physical colors of color-bar stimuli (i.e., green, yellow, orange, and purple with RGB values of [0 170 0], [173 145 0], [220 120 0], and [230 0 255], respectively; iso-luminace of 14.10 cd/m^2^ ± 0.72 SD) with four buttons on a numeric keypad (‘7’, ‘4’, ‘1’, and ‘0’), which they pressed using their index, middle, ring, and little fingers of their right hand, respectively. Each trial began with the presentation of the color-bar stimulus (size = 4.30° x 21.70° visual angle), which appeared 2.39° visual angle below a central black fixation dot (radius = 0.38° visual angle). Each color-bar stimulus was presented for 1000ms, and participants were instructed to report its physical color as quickly and accurately as possible before the stimulus disappeared. 300ms following the stimulus offset, subjects received feedback on their performance for that trial (‘C’ for correct responses, ‘I’ for incorrect responses, and ‘M’ for misses) for 200ms. The inter-trial interval was randomly drawn from the uniform distribution of 500-1500ms. Each subject completed one block of the stimulus-response mapping task, which consisted of 144 trials in total and lasted approximately 6 minutes (36 trials per each color; trial order was pseudo-randomized).

Immediately after completing the stimulus-response mapping task, subjects performed an adapted version of the color-naming Stroop task (Figure 1a). They were instructed to fixate at a central fixation point while attending to the color-bar stimulus and ignoring the letter-string stimulus (all letters were capitalized; font type = ‘Arial’; font size = 3.34o visual angle in height), which appeared over of the color-bar stimulus. The letter-string stimulus could be a non-word (i.e., neutral; e.g., color-letter = purple-AEGNRL) or a word that was semantically congruent (e.g., color-letter = purple-PURPLE) or incongruent (e.g., color-letter = purple-GREEN) with respect to the physical color of the color-bar stimulus. To concurrently monitor sensory responses evoked by the color-bar and letter-string stimuli, the two stimuli were flickered at different frequencies for 1500ms (20Hz color-bar and 30Hz letter-string or 30Hz color-bar and 20Hz letter-string; the frequency assignments were counterbalanced block-by-block). This stimulus-frequency-tagging technique allowed us to obtain steady-state visually evoked potentials (SSVEPs) elicited by the color-bar and letter-string stimuli (relevant and irrelevant stimuli, respectively). The flicker frequencies of 20Hz and 30Hz were chosen based on previously established methods in order to restrict SSVEP measurements to entrained activity in the visual cortex, and to avoid spectral overlap with intrinsic theta (4-7Hz) and alpha (8-14Hz) oscillations (e.g., Müller et al., 1998; O’Connell et al., 2012; Bridwell et al., 2013; Garcia et al., 2013; Itthipuripat et al., 2013a, 2014b). Participants were instructed to report the physical color of the color-bar stimulus as quickly and accurately as possible. The size, luminance and RGB values of the color-bar stimuli, feedback duration, and ITI were identical to those used in the stimulus-response mapping task. Subjects completed four blocks of the Stroop task where color bar and letter-string stimuli were flashed at 20Hz and 30Hz and four blocks where they were flashed at 30Hz and 20 Hz (the order of block types were counterbalanced across subjects). Each block contained 144 trials (48 congruent trials, 48 neutral trials, and 48 incongruent trials), and each block lasted about 7.2 minutes.

### Statistical Analysis of Behavioral Data

For each subject, we computed hit rates, mean response times on correct trials (correct RTs), and miss rates on congruent, neutral, and incongruent trials separately for the blocks that had different frequency assignments to the color-bar and letter-string stimuli. The within-subject SEM of the data was calculated by removing the mean value of each congruency condition and each frequency assignment from the individual subject data before computing the SEM (Loftus and Masson, 1994). Then, we performed repeated-measures ANOVAs with within-subject factors of congruency and frequency assignment to test the main effects of these two factors on hit rates, correct RTs, and miss rates. Post-hoc t-tests (2-tailed) were then used to test differences in hit rates, correct RTs, and miss rates between the congruent and incongruent conditions, between the congruent and neutral conditions, and between the neutral and incongruent conditions, respectively. We used the false discovery rate (FDR) method to correct for multiple comparisons with the corrected threshold of 0.05 (Benjamini and Hochberg, 1995).

### EEG Data Acquisition

EEG data were recorded with a 64+8 electrode Biosemi ActiveTwo system (Biosemi Instrumentation) using a sampling rate of 512 Hz. Two reference electrodes were placed on the left and right mastoids. Blinks and vertical eye movements were monitored using four external electrodes affixed above and below the eyes. Horizontal eye movements were monitored using another pair of external electrodes affixed near the outer canthi of the left and right eyes. The EEG data were referenced on-line to the CMS-DRL electrode and the data offsets in all electrodes were maintained below 20uV (a standard criterion for this active electrode system).

### EEG Data Preprocessing and Analysis

EEG data were preprocessed using a combination of EEGLab11.0.3.1b (Delorme and Makeig, 2004) and custom MATLAB scripts. The continuous EEG data were first re-referenced to the algebraic mean of the left and right mastoid electrodes, then filtered by applying 0.25-Hz high-pass and 55-Hz low-pass Butterworth filters (3rd order). Next, the continuous EEG data were segmented into epochs extending from 1000ms before to 2500ms after trial onset. Independent component analysis (ICA) was then applied in order to remove prominent eye blinks (Makeig et al., 1996). Trials containing residual eye movements, muscle activity, drifts, and other artifacts were removed using threshold rejection and visual inspection, which resulted in the removal of 7.79% ± 8.87 SD of trials across all 30 subjects.

Next, we wavelet-filtered the artifact-free EEG data using Gaussian filters centered at 4-7Hz (1-Hz steps), 8-13Hz (1-Hz steps), 20Hz, and 30Hz with a fractional bandwidth of 0.2. This method yielded analytic coefficient values of the EEG data from across these specific frequency bands (see similar methods in Canolty et al., 2007; Roach and Mathalon, 2008; Itthipuripat et al., 2013b; Freeman et al., 2016). To compute SSVEPs evoked by color-bar and letter-string stimuli, the analytic coefficient values at the driving frequencies of 20Hz and 30 Hz for individual trials were sorted by the driving stimuli (the color-bar and letter-string stimuli) that were semantically congruent, neutral, or incongruent. For each congruency condition, the data from correct trials were also sorted by RTs into 10 bins (from fast to slow RTs in incrementing 10%-percentile steps). The sorted coefficient values were then averaged across trials and the absolute values of these averaged coefficients were then computed, yielding the analytic amplitude of SSVEP signals. Then, we baseline-corrected SSVEP amplitude across time for each condition by subtracting the mean amplitude 500-0ms before the stimulus onset. To obtain the amplitude of induced theta and alpha oscillations in the EEG data, we computed the absolute values of the coefficient values on a trial-by-trial basis from 4-7Hz and from 8-13Hz, respectively. Next, the single-trial data were sorted based on congruency and RT, averaged across trials in each experimental bin, and baseline-corrected from 500-0ms before the stimulus onset. The within-subject SEM of the SSVEP, theta, and alpha data was calculated by removing the mean value of each congruency condition and each RT bin from the individual subject data before computing the SEM (Loftus and Masson, 1994).

### Statistical Analysis of EEG Data

For statistical evaluation of the data, we used repeated-measures ANOVAs to test the main effects of congruency and RT and their interaction on target-related SSVEPs (i.e., SSVEPs elicited by color-bar stimuli), distractor-related SSVEPs (i.e., SSVEPs elicited by letter-string stimuli), induced frontal theta activity, and induced posterior alpha activity from 500ms before to 1500ms after stimulus onset. Because we displayed the stimuli at the center of the screen, the SSVEP, theta, and alpha data were obtained from the central occipital electrodes (O1, Oz, and O2), central frontal electrodes (F1, Fz, and F2), and posterior-occipital electrodes (PO1, POz, and PO2), respectively. These electrodes have also been used as standard electrodes of interest to analyze SSVEP, theta, and alpha in previous studies (Sauseng et al., 2005; Kelly et al., 2006, 2009; Andersen and Muller, 2010; Cavanagh et al., 2011, 2012; Foxe and Snyder, 2011; Andersen et al., 2012; Störmer et al., 2013; Itthipuripat et al., 2013b; Freeman et al., 2016).

For any consecutive time points where ANOVAs showed a significant main effect of congruency, we averaged the data across the significant time points, and performed post-hoc t-tests (2-tailed) to determine whether the main effects were driven by differences between the congruent and incongruent conditions, between the congruent and neutral conditions, and/or between the neutral and incongruent conditions. For any consecutive time points where ANOVAs showed significant main effects of RT, we averaged the data across those time points and used one-way repeated-measured ANOVAs with a within-subject factor of RT to test whether effects were consistent across the congruent, neutral, and incongruent conditions. For each statistical evaluation that was performed separately on the target-related SSVEPs, the distractor-related SSVEPs, the frontal theta activity, and the posterior alpha activity, multiple comparisons across time points were FDR-corrected with a corrected threshold of 0.05 (Benjamini and Hochberg, 1995). For all post-hoc analyses, multiple comparisons were also FDR-corrected using the same threshold of 0.05.

## Results

### Behavioral results

Consistent with many previous studies employing variants of the Stroop task, incongruent color-word pairings led to significant effects on hit rates, correct RTs, and miss rates (F(2, 58)’s = 23.91, 131.61, and 8.53, respectively, with all p’s < 0.001) (Figures 1b-s) (Jensen and Rohwer JR., 1966; Stroop, 1992; Liotti et al., 2000; West and Alain, 2000; Zysset et al., 2001; Kane and Engle, 2003; Atkinson et al., 2003; Hanslmayr et al., 2008; Appelbaum et al., 2009, 2012; Huster et al., 2009; Krebs et al., 2010, 2013; Coderre et al., 2011; Caldas et al., 2012; Donohue et al., 2013, 2016; van den Berg et al., 2014). Post-hoc t-tests revealed that hit rates in the incongruent condition were significantly lower than hit rates in the neutral and congruent conditions (t(29)’s = 5.84 and 4.85, respectively, both p’s < 0.001, FDR-corrected), with no significant difference between the congruent and neutral conditions (t(29) = 0.97, p = 0.340). Correct RTs in the incongruent condition were significantly longer than correct RTs in the neutral and congruent conditions (t(29)’s = 10.14 and 12.89, respectively, with both p’s < 0.001, FDR-corrected). Correct RTs in the neutral condition were also significantly longer than correct RTs in the congruent condition (t(29) = 8.55, p < 0.001, FDR-corrected). Miss rates in the incongruent condition were also higher than miss rates in the neutral and congruent conditions (t(29)’s = 3.14 and 3.00, respectively, with both p’s < 0.01, FDR-corrected) with no difference between the neutral and congruent conditions (t(29) = 0.55, p = 0.587). In addition, there were no differences in hit rates, correct RTs, or miss rates across trials that contained a 20-Hz flickering color-bar and 30-Hz flickering letter-string and trials that contained 30-Hz flickering color-bar and 20-Hz flickering letter-string (F(1, 29)’s = 0.03, 0.10, 1.80, p’s = 0.87, 0.76, and 0.19, respectively).

### EEG results

#### Target-related and distractor-related SSVEPs

Overall SSVEP results are not consistent with the early selective sensory processing account. There were robust target-related SSVEPs evoked by task-relevant color-bar stimuli over occipital electrodes (Figure 2a). However, the amplitude of the SSVEPs was not modulated by congruency (F(2, 58)’s ≤ 4.50, p’s ≥ 0.015, n.s. FDR-corrected) or RT (F(9, 251)’s ≤ 1.96, p’s ≥ 0.044, n.s. FDR-corrected), and there was no interaction between the two factors (F(18, 522)’s ≤ 1.50, p’s ≥ 0.083, n.s. FDR-corrected). Similarly, we observed robust distractor-related SSVEPs evoked by task-irrelevant letter-string stimuli but there was no effect of congruency on the amplitude of the response (F(2, 58)’s ≤ 3.41, p’s ≥ 0.039, n.s. FDR-corrected) (Figure 2b-c). However, the distractor-related SSVEP amplitude rose just before and fell right after motor responses were executed, resulting in significant main effects of RT from 295-385ms and 723-1102ms post-stimulus (F(9, 251)’s ≥ 2.27, p’s ≤ 0.018, FDR-corrected). More specifically, the amplitude of word-SSVEPs was higher for fast compared to slow trials from 295-385ms post-stimulus but word-SSVEPs were slower for fast compared to slow trials from 723-1102ms post-stimulus. This general result was observed in all congruency conditions: congruent (early window: F(1, 29)= 40.27, p < 0.001; late window: F(1, 29)= 49.72, p < 0.001), neutral (early window: F(1, 29)= 44.04, p < 0.001; late window: F(1, 29)= 57.70, p < 0.001), and incongruent (early window: F(1, 29)= 43.12, p < 0.001; late window: F(1, 29)=54.09, p < 0.001, all tests were FDR-corrected). In addition, there was no interaction between congruency and RT on the distractor-related SSVEP amplitude at any time point (F(18, 522)’s ≤ 1.52, p’s ≥ 0.079, n.s. FDR-corrected).

**Figure 2.**
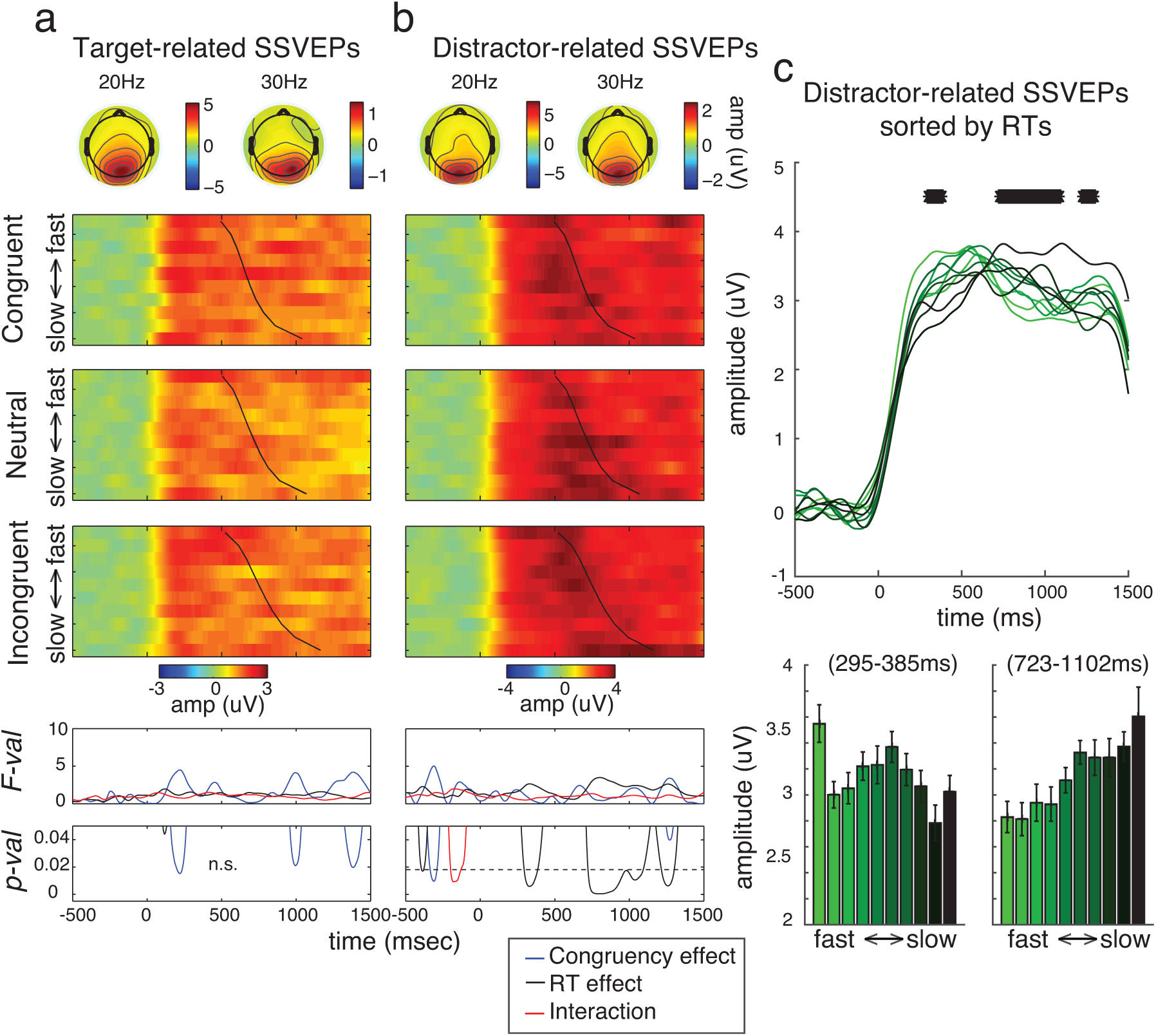
(a) SSVEPs evoked by task-relevant color-bar stimuli (target-related SSVEPs) and (b) task-irrelevant letter-string stimuli (distractor-related SSVEPs). The data were sorted by congruency conditions and RTs. There were no main effects of congruency or RTs on color-SSVEPs, and no interaction between the two factors on target-related SSVEPs. On the other hand, while there was no main effect of congruency or interaction, there were significant main effects of RT on distractor-related SSVEPs. The faster the distractor-related SSVEPs increased in amplitude, the faster correct responses were executed. This finding is inconsistent with the prediction of the selective sensory modulation account. A black dotted line in the last panel of (b) shows a significant FDR-corrected threshold of 0.018 for RT effects on distractor-related SSVEPs. Black lines in the heat maps in (a-b) represent mean RTs for individual RT bins. Topographical maps above the heat maps show SSVEP amplitude averaged across the entire stimulus duration and collapsed across congruency conditions for the color-bar and letter-string stimuli of 20Hz and 30Hz. (c) Distractor-related SSVEPs sorted by RTs, collapsed across congruency conditions for all time points (top) and for each of the significant windows (bottom). Light-to-dark colors correspond fast-to-slow trials. Black * signs in (c) show significant main effects of RT (FDR-corrected). Error bars represent ±1 within-subject S.E.M.

#### Frontal theta activity

There were significant congruency effects on the amplitude of induced theta activity recorded from the frontal electrodes from 484-604ms after stimulus onset (F(2, 58)’s ≥ 5.22, p’s ≤ 0.008, FDR-corrected) (Figures 3a-b). Consistent with previous studies, the theta amplitude in the incongruent condition was significantly higher than the neutral condition and the congruent condition (t(29) = 2.89, p =0.007 and t(29) = 2.51, p =0.018, respectively, FDR-corrected) (e.g., Hanslmayr et al., 2008; Ergen et al., 2014), without any difference between the congruent and neutral conditions (t(29) = 0.41, p =0.683). Moreover, we observed that the rising and falling time course of frontal theta activity closely matched RTs (significant main effects of RT from 273-430ms and 533-1500ms post-stimulus; F(9, 251)’s ≥ 2.03, p’s ≤ 0.037, FDR-corrected) (Figures 3a&c). Specifically, we observed higher theta amplitude for fast compared to slow trials in the early time window but higher theta amplitude for slow compared to fast trials in the later time window. Moreover, this result was consistent across all congruency conditions: congruent (early window: F(1, 29)= 18.17, p < 0.001; late window: F(1, 29)= 5.14, p < 0.001), neutral (early window: F(1, 29)= 21.65, p < 0.001; late window: F(1, 29)= 4.86, p < 0.001), and incongruent (early window: F(1, 29)= 19.92, p < 0.001; late window: F(1, 29)= 7.47, p < 0.001). Finally, there was no interaction between congruency and RT on theta amplitude (F(18, 522)’s ≤ 1.69, p’s ≥ 0.037, n.s. FDR-corrected).

**Figure 3.**
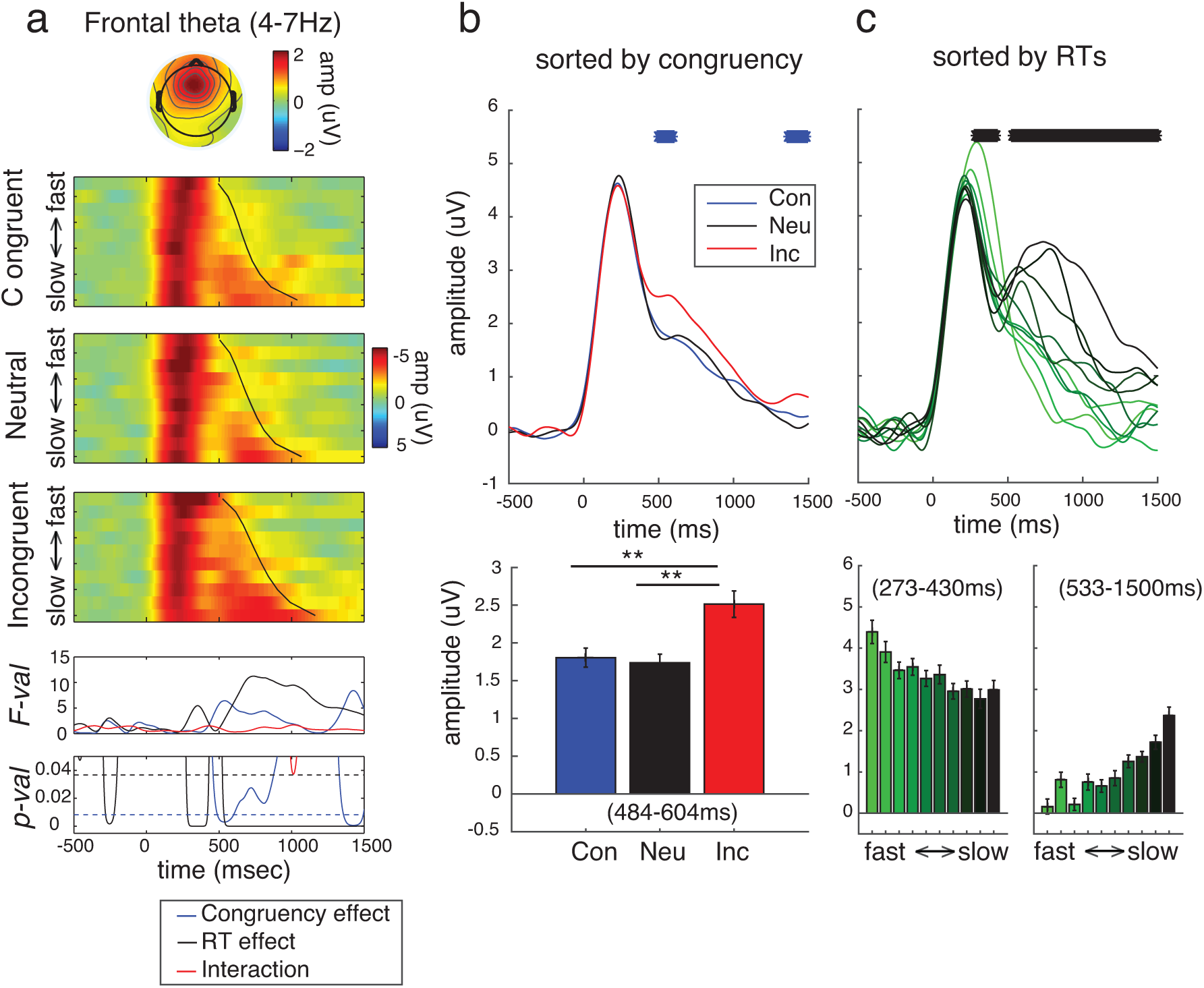
(a) Stimulus-locked frontal theta activity sorted by congruency conditions and RTs. A topographical map above the heat maps shows theta amplitude averaged across the entire stimulus duration and collapsed across congruency conditions. Black lines in the heat maps represent mean RTs for individual RT bins. Blue and black dotted lines in the last panel of (a) show significant FDR-corrected thresholds of 0.008 and 0.037 for congruency and RT effects, respectively. (b) Frontal theta activity increased for the incongruent compared to the neutral and congruent trials. (c) The higher the frontal theta activity was at the early time window 273-430ms, the faster correct responses were made. For faster trials, the theta amplitude also dropped faster leading to significant main effects of RT in the later time window of 533-1500ms. Blue and black * signs in the top panels of (b-c) show significant main effects of congruency and RT (FDR-corrected), respectively. ** in the bottom panel of (b) show significant pair-wise difference between congruency conditions with p’s < 0.01. Light-to-dark colors in (c) correspond fast-to-slow trials. Error bars represent within subject S.E.M.

#### Posterior alpha activity

There were significant congruency effects on the amplitude of stimulus-locked alpha at the posterior-occipital electrodes from 510-1211ms after stimulus onset (F(2, 58)’s ≥ 3.86, p’s ≤ 0.027, FDR-corrected) (Figures 4a-b). Specifically, alpha amplitude decreased less in the incongruent condition than in the neutral (t(29) = 4.64, p <0.001) and congruent conditions (t(29) = 3.51, p =0.002). This is consistent with the observation that correct RTs in the incongruent condition were longer and with the proposal that events with high levels of cognitive conflict (i.e., incongruent trials) may lead to more post-conflict attention and task engagement (Talsma et al., 2010; Zimmer et al., 2010). Collapsed across congruency conditions, we also observed a more sustained decrease in alpha amplitude, from 469-1402ms for slower compared to faster trials (F(9, 251)’s ≥ 2.09, p’s ≤ 0.031, FDR-corrected) (Figures 4a&c). This sustained decrease in alpha amplitude was also observed in each condition considered separately: congruent (F(1, 29)= 71.70, p < 0.001), neutral (F(1, 29)= 63.98, p < 0.001), and incongruent conditions (F(1, 29)= 99.69, p < 0.001). Last, there was no interaction between congruency and RT on alpha amplitude (F(18, 522) ≤ 1.80, p ≥ 0.022, n.s. FDR-corrected).

**Figure 4.**
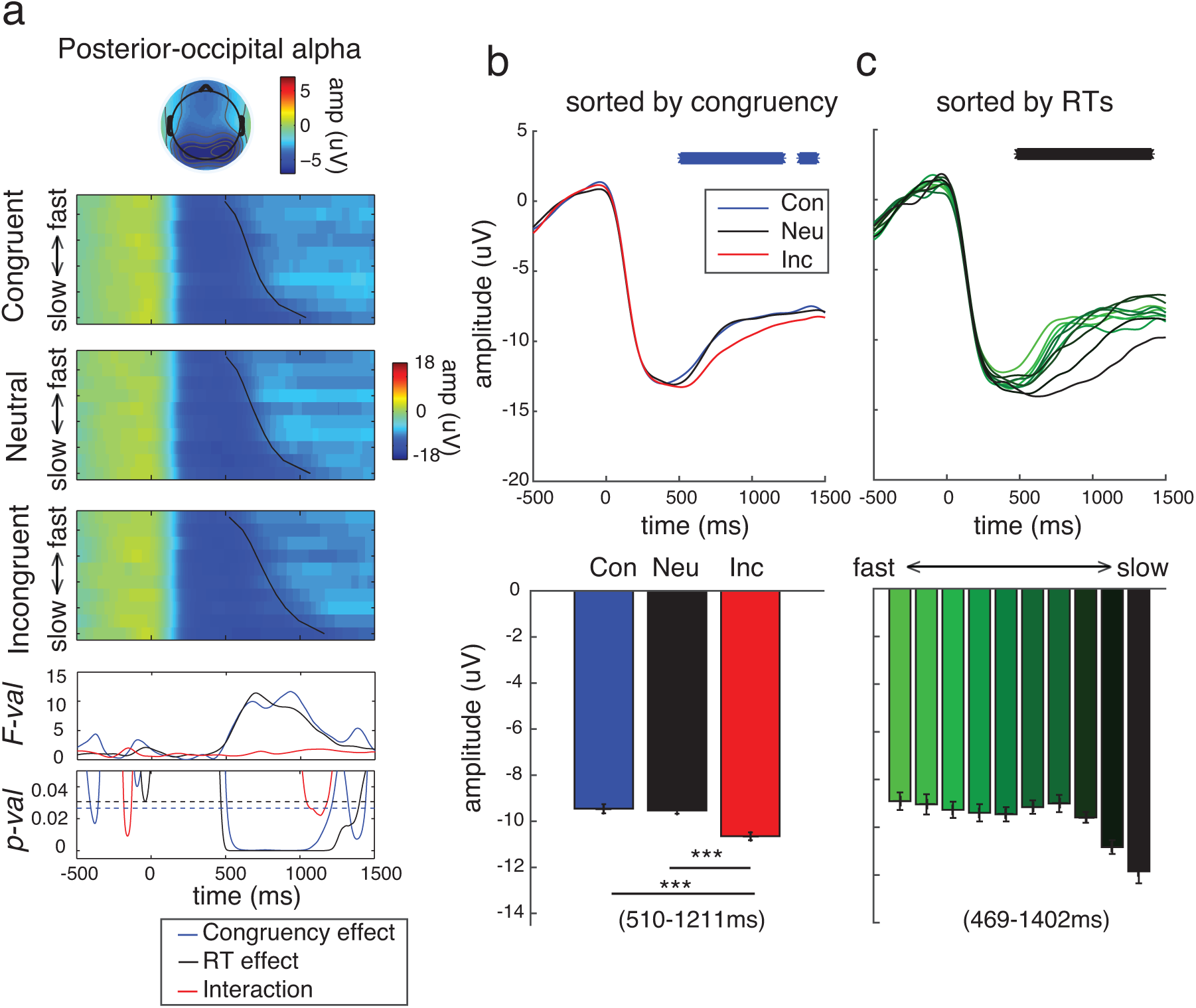
(a) Stimulus-locked posterior-occipital alpha activity sorted by congruency conditions and RTs. A topographical map above the heat maps shows alpha amplitude averaged across the entire stimulus duration and collapsed across congruency conditions. Black lines in the heat maps represent mean RTs for individual RT bins. Blue and black dotted lines in the last panel of (a) show significant FDR-corrected thresholds of 0.027 and 0.031 for congruency and RT effects, respectively. (b) Post-stimulus alpha activity decreased for the incongruent compared to the neutral and congruent trials. (c) The longer post-stimulus alpha reduction sustained, the longer correct responses were made. Blue and black * signs in the top panels of (b-c) show significant main effects of congruency and RT (FDR-corrected), respectively. *** in the bottom panel of (b) show significant pair-wise difference between congruency conditions with p’s < 0.001. Light-to-dark colors in (c) correspond fast-to-slow trials. Error bars represent with-in subject S.E.M.

#### Auxiliary EEG results

It has been proposed that the subthalamic nucleus (STN), a subcortical structure in the basal ganglia, communicates with the prefrontal cortex via theta frequency oscillations to facilitate cognitive control (Cavanagh et al., 2011; Itthipuripat et al., 2013b; Zavala et al., 2013, 2014, 2015; Cavanagh and Frank, 2014). A recent study recorded theta activity directly from the STN in human participants performing the Eriksen flanker task and found that there were significant differences in STN theta activity between slow incongruent and all congruent trials. However, no difference was observed between fast incongruent and all congruent trials (Zavala et al., 2013). The authors suggested that this pattern might be observed because distractors (i.e., flankers) were successfully ignored on fast incongruent trials, consistent with the selective sensory modulation account (Zavala et al., 2013). Here, we explored our data to determine if a similar analysis of frontal theta activity would yield a pattern similar to that observed in the STN and whether these results could be explained by the selective sensory modulation hypothesis as suggested by Zavala et al. (2013). Similar to the STN theta results (Zavala et al., 2013), we observed significant increases in frontal theta activity on slow incongruent compared to fast incongruent and all congruent trials from 566-1463ms and from 494-1500ms, respectively (|t(29)|’s ≥ 2.44, p’s ≤ 0.021, FDR-corrected) (Figure 5a). However, there was no difference in frontal theta activity between fast incongruent and all congruent trails (|t(29)|’s ≤ 2.10, p’s ≥ 0.044, n.s. FDR-corrected). These results follow a time course similar to that observed for the posterior-occipital alpha activity where there was a significant decrease in alpha amplitude on slow incongruent compared to fast incongruent and to all congruent trials from 625-1062ms and 500-1172ms post-stimulus, respectively (|t(29)|’s ≥ 2.67, p ≤ 0.012, FDR-corrected) (Figure 5b). However, there was no significant difference between fast incongruent and congruent trials (|t(29)|’s ≤ 2.06, p’s ≥ 0.049, n.s. FDR-corrected). In contrast to the theta and alpha results, we found no significant difference in target-related and distractor-related SSVEPs across congruent, fast incongruent, and slow incongruent trials (|t(29)|’s ≤ 2.37, p’s ≥ 0.025, n.s. FDR-corrected) (Figures 5c-d).

**Figure 5.**
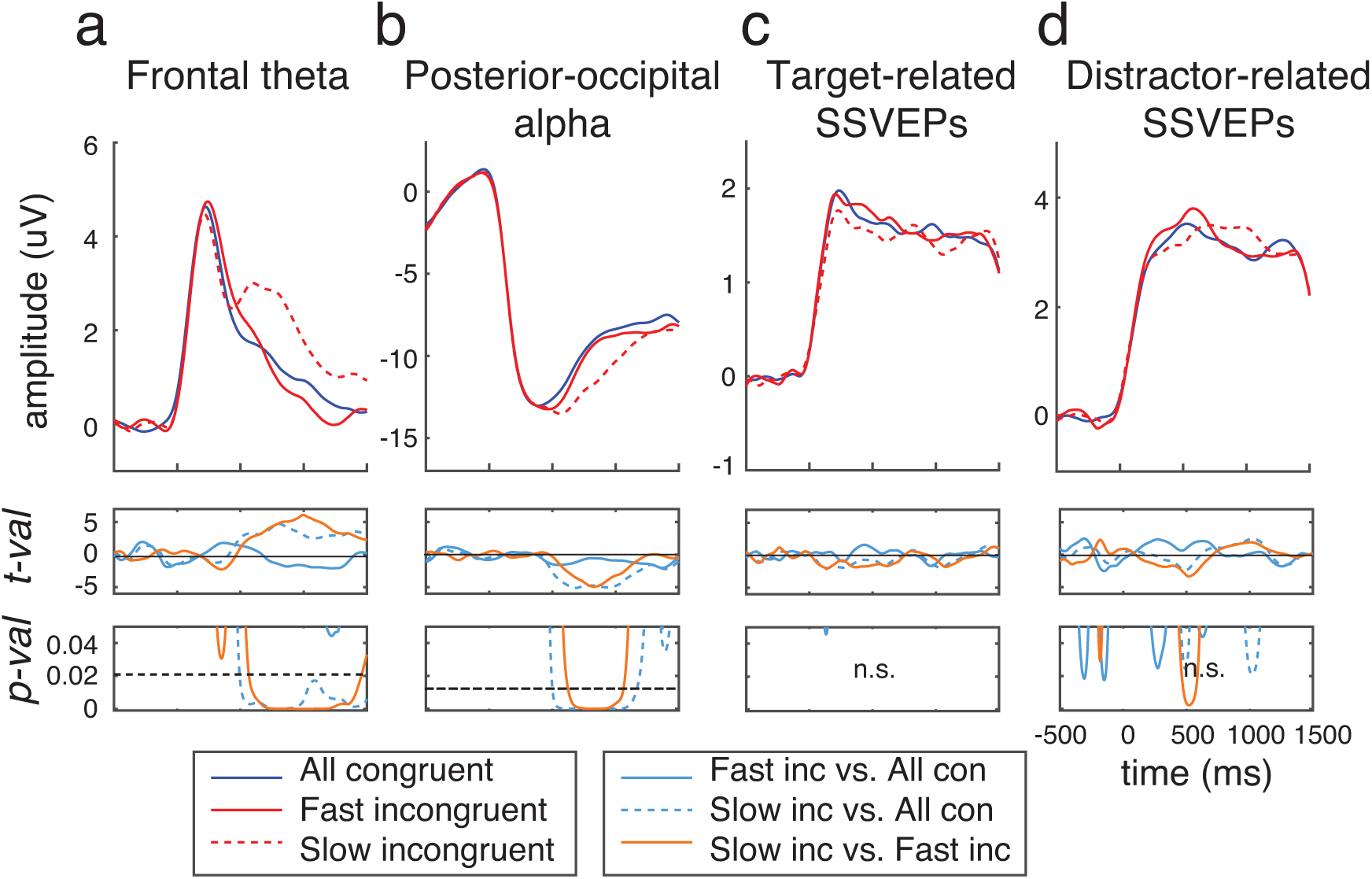
Comparisons of (a) frontal theta activity, (b) posterior-occipital alpha activity, (c) target-related SSVEPs, and (d) distractor-related SSVEPs across all congruent trials, fast incongruent trials, and slow incongruent trials. Dotted black lines in the last panels of (a-b) represent significant FDR-corrected thresholds of 0.021 and 0.012, respectively.

## Discussion

The present study evaluated the relative contributions of selective sensory modulations and frontal executive functions on the efficiency of decision-making in the face of cognitive conflict. We used a novel version of the Stroop task, where sensory signals (i.e., SSVEPs) evoked by relevant color-bar and irrelevant letter-string stimuli were tagged using two different stimulus-flicker frequencies. Counter to the ‘selective sensory modulation’ account, we found no changes in target-related SSVEPs either as a function of congruency or as a function of RT. Moreover, distractor-related SSVEPs appeared to change as a function of RT but in the opposite direction as predicted by this account. Specifically, we found that higher distractor-related SSVEP amplitude as early as 295ms post-stimulus predicted faster RTs, irrespective of congruency condition. This result supports the idea that the processing of letter-string stimuli (i.e., distractors) in the Stroop task is automatic and was not disengaged until a decision about the color-bar stimulus was made. In contrast, we found that higher frontal theta amplitude 273-430ms post-stimulus, which is in a similar time range as the significant RT effects on distractor-related SSVEPs (295-385ms), predicted faster RTs. Overall, the data suggest that decision-making efficiency under high-order cognitive conflict relies primarily on the activation state of the frontal executive control network, rather than low-level selective sensory modulations.

Note that higher frontal theta activity in an early time window (273-430ms) predicted faster RTs irrespective of stimulus congruency. In addition, theta was high on incongruent trials from 484-604ms, which were associated with the longest RTs. This may seem difficult to reconcile at first glance, but the effects of congruency on theta activity occurred in a later time window (484-604ms) compared to effects of general RT changes on theta activity (273-430ms). These bidirectional modulations of theta oscillations at different temporal windows suggest that the increase in frontal theta activity indexes at least two cognitive operations occurring at different points in time. In the early time window, frontal theta activity may serve a more general executive control function, such as the initiation of executive control mechanisms engaged by the stimulus and task, irrespective of conflict level (Cavanagh et al., 2012; Cavanagh and Frank, 2014). On the other hand, in the later time window, the increase in theta activity on the relatively slow incongruent trials may reflect additional frontal activity that is recruited to support conflict-monitoring functions (Carter et al., 1998; Botvinick et al., 1999, 2001, 2004; Cavanagh et al., 2012; Cavanagh and Frank, 2014; Zavala et al., 2014). Thus, the present results are consistent with studies showing that frontal theta activity indexes multiple attributes of executive functions including conflict-monitoring, error detection, response inhibition, novelty detection, and working memory (D’Esposito et al., 1995; Carter et al., 1998; Botvinick et al., 1999, 2001, 2004; Kane and Engle, 2003; Curtis and Esposito, 2003; Aron et al., 2004, 2014; Ridderinkhof et al., 2004; Cavanagh et al., 2011, 2012; Itthipuripat et al., 2013b; Cavanagh and Frank, 2014; Wessel and Aron, 2017).

Interestingly, while the auxiliary EEG analysis showed a congruency effect on frontal theta activity when comparing slow incongruent and all congruent trials, there was no significant difference in theta activity between fast congruent and all congruent conditions when RTs were matched. Similar results were also reported in a recent study that measured theta activity of the STN in humans performing the Eriksen flanker task (Zavala et al., 2013). The similarity of the results recorded at the scalp over frontal cortex and modulations observed directly in the STN suggests that areas of frontal cortex and the basal ganglia may rely on communication in the theta band to support their putative functions in cognitive control and cognitive interference tasks (Cavanagh et al., 2011; Itthipuripat et al., 2013b; Zavala et al., 2013, 2014, 2015; Cavanagh and Frank, 2014). Zavala and colleagues (2013) reasoned that the absence of differences in the STN theta activity and RTs between fast incongruent trials and all congruent trials was due to the fact that on fast incongruent trials subjects were able to successfully suppress irrelevant sensory signals, thus reducing conflict generated by the distractors. However, in the present study, we directly measured sensory signals evoked by the distractors and found that this was not the case. Our results are thus consistent with the idea that the speed and efficiency of decision making during higher-order cognitive conflict relies primarily on interactions between the frontal executive control network and sub-regions of the basal ganglia rather than low-level selective sensory modulations.

In addition to SSVEPs and frontal theta activity, we also examined the effect of congruency and RTs on post-stimulus posterior-occipital alpha activity, which has been previously used as an index of attentional control and general task engagement (von Stein et al., 2000; Fries, 2001; Sauseng et al., 2005; Klimesch et al., 2007; Rihs et al., 2007; Fries et al., 2008; Hanslmayr et al., 2008; Busch et al., 2009; Kelly et al., 2009; Mathewson et al., 2009; Zhang et al., 2010; Foxe and Snyder, 2011; Bosman et al., 2012). Note that topographical patterns of reductions in posterior alpha activity have also been shown to track the active maintenance of spatial attention and visual short-term spatial memory (VSTM) in a selective manner (Sauseng et al., 2005; Kelly et al., 2006; Foxe and Snyder, 2011; Bosman et al., 2012; Foster et al., 2016, 2017; Samaha et al., 2016). However, the null effect of SSVEP modulations in the present study suggests that the temporally extended reduction of alpha activity on slower trials reflects more sustained vigilance and task engagement rather than selective attention to the relevant sensory stimulus. In line with previous studies proposing that cognitive conflict could lead to more sustained attention and task engagement (Talsma et al., 2010; Zimmer et al., 2010), we observed a higher degree of post-stimulus alpha reduction for incongruent compared to neutral and congruent trials. In addition, higher amplitude post-stimulus alpha reductions were observed in a similar time window when RTs were longer, irrespective of congruence. Similar to frontal theta, we found no difference between fast incongruent and all congruent trials when RTs were matched, but a higher degree of alpha reduction for slow incongruent trials compared to slow incongruent and all congruent trials. The similarity of modulations of frontal theta and posterior alpha activity suggests a tight coupling between prefrontal and frontoparietal networks during the resolution of conflicting stimulus inputs.

## Acknowledgement

We thank Greg L. Appelbaum for useful comments and suggestions. Funding provided by NIH R01-EY025872 grant to JTS, a James S. McDonnell Foundation Scholar Award to JTS, a Howard Hughes Medical Institute International Student Research Fellowship to SI and a Royal Thai Scholarship from the Ministry of Science and Technology in Thailand to SI.

